# Light-dependent decomposition of FICZ, an endogenous ligand of the aryl hydrocarbon receptor

**DOI:** 10.1101/158980

**Authors:** Cunyu Zhang, Katrina L. Creech, William J. Zuercher, Timothy M. Willson

## Abstract

An efficient and scalable synthesis of 6-formylindolo[3,2-b]carbazole (FICZ) has been developed to provide large quantities of this physiologically important ligand of the aryl hydrocarbon receptor. Photo-decomposition of FICZ revealed a new non-enzymatic light-assisted mechanism for its conversion to a biologically less active quinone. The light-dependent synthesis and decomposition of FICZ makes it a candidate hormone to link sun exposure to regulation of biological pathways in peripheral tissues.

## Introduction

The day/night cycle controls basic aspects of human physiology, such as when we sleep and core body temperature. In mammals, light detected by photoreceptors in the eye entrains the rhythm of a molecular clock within specialized cells of the hypothalamus. Neuronal signaling and timed release of hormones, such as corticosterone and melatonin, are used transmit these circadian rhythms to other cells [1]. Cyclical secretion of the hormone melatonin by the pineal gland is a primary regulator of sleep and wakefulness. Melatonin has also emerged as one of the most biologically relevant molecules with regards to mental health and wellbeing suggesting an important link between small molecules and the day/night cycle [2]. The biology of peripheral organs also demonstrates important day/night differences, but less is known about how these are controlled by secreted hormones or how they are synchronized with the central clock. In bacteria, protein components of the molecular clock can be directly regulated by light [3], but in humans no related processes are known for photochemical regulation of peripheral cell signaling.

6-Formylindolo[3,2-b]carbazole (FICZ, 1) is an interesting photoproduct of tryptophan (Fig 1) that was isolated by the Rannug laboratory as a light-induced ligand for the aryl hydrocarbon receptor (AhR) [4]. 1 is the most potent naturally-occurring AhR agonist. Strong evidence for its formation in humans by the action of light on the skin and its metabolism by the AhR-inducible CYP1A have been presented [5–7]. As such, 1 fulfills many of the requirements for a hormone that links sunlight to regulation of peripheral biological rhythms [8]. Remarkably, for a potential light-regulated hormone, relatively little is known about its photo-stability or potential for photo-degradation (Fig 1).

**Fig 1.**
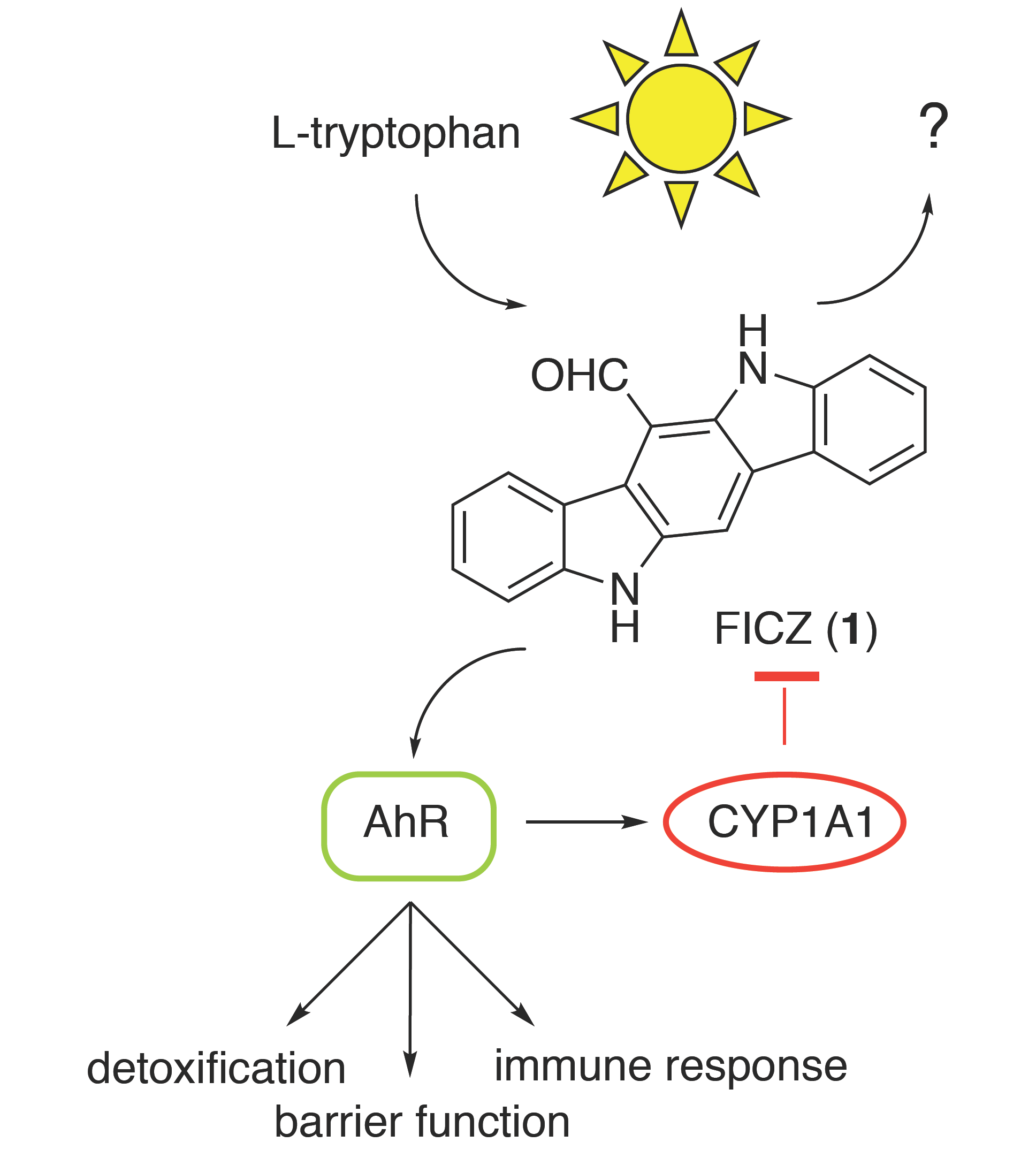
Light dependent synthesis of FICZ. 6-Formylindolo[3,2-b]carbazole (FICZ, **1)**is formed by the action of light on tryptophan. FICZ activation of AhR regulates important physiological pathways and also induces expression of CYP1A1 as a feedback mechanism to induce its own metabolism.

Despite the importance of **1** as a reference compound for AhR research, it is available in only small quantities from commercial vendors. The chemical synthesis of **1** was first reported by Bergman through a low yielding route that required purification of the final product by sublimation under reduced pressure [9]. Subsequent improvements [10, 11] were made to facilitate the synthesis of related carbazoles, but no simple scalable route for the production of **1** has been described. A key challenge in the synthesis has been the difficulty in handling the intermediate carbazoles due to their poor solubility. Even the most recent syntheses were completed on small scales that yield only a few milligrams of product [12, 13]. To address these issues, we developed an optimized synthesis that allows production of **1** in gram quantities for the first time and investigated the photochemical stability of the molecule.

## Materials and methods

Starting materials, reagents, and solvents were obtained from commercial sources and used as received. Indolo[3,2-b]carbazole was purchased from AstaTech. Intestinal human colon adenocarcinoma cells LS180 stably transfected with a β-lactamase reporter gene downstream of the CYP1A1 promoter were obtained from Invitrogen.

^1^H NMR and proton decoupled ^13^C NMR spectra were recorded on a Bruker 400-MHz spectrometer in CDCl_3_ or DMSO-*d*_6_ with TMS as the internal reference. Chemical shifts (δ) are reported in parts per million (ppm) relative to deuterated solvent as the internal standard (δH: CDCl_3_ 7.26 ppm, DMSO-*d*_6_ 2.50 ppm), coupling constants (J) are in hertz (Hz). Peak multiplicities are expressed as follows: singlet (s), doublet (d), doublet of doublets (dd), triplet (t), quartet (q), multiplet (m), and broad singlet (br s). Melting points were recorded with a Büchi 520 apparatus and are uncorrected. Purification of intermediates and final products, if mentioned, was carried out on normal phase using an ISCO CombiFlash system and prepacked SiO_2_ cartridges eluted with optimized gradients of heptane-ethyl acetate mixture. Progress of the reactions was monitored by thin-layer chromatography (TLC) analysis (Merck, 0.2 mm silica gel 60 F254 on glass plates) or by LC-MS (described below). The purity of all products was established on a LCMS method:

LC conditions: UPLC analysis was conducted on a Phenomenex Kinetex 1.7 um XB-C18 column at 40 °C. 0.2 uL of sample was injected using a partial loop with needle overfill injection mode. The gradient employed was: mobile phase A - water + 0.2% v/v formic acid., mobile phase B - acetonitrile + 0.15% v/v formic acid. UV detection was provided by summed absorbance signal from 210 to 350 nm scanning at 40Hz. MS detection by Waters SQD LBA555 using alternating positive/negative electrospray with scan range = 125-1200 amu, scan time =110 msec, and interscan delay = 50 msec.

### (1-(Phenylsulfonyl)-1H-indol-2-yl)(1-(phenylsulfonyl)-1H-indol-3-yl)methanol

To a stirred solution of 1-(phenylsulfonyl)-1H-indole (9.02 g, 35.0 mmol) in dry 2-methyltetrahydrofuran (200 mL) at −78 °C was added 1.6 M *n*-butyllithium in hexane (28 mL, 44.8 mmol) slowly over 15min. The mixture was warmed to room temperature and stirred for 0.5-1 h. The reaction mixture became a slurry. It was cooled back to −78 °C and a solution of 1-(phenylsulfonyl)1-(phenylsulfonyl)-1H-indole-3-carbaldehyde (10 g, 35 mmol) in 110 ml of THF was added dropwise over 20 min. The reaction mixture became a homogeneous solution. After stirring for 30 min with continued cooling by a dry ice-acetone bath, LCMS revealed the aldehyde had been consumed. The reaction was quenched by slowly adding 1N HCl aq. solution (100 mL) and then allowed to warm to room temperature and stirred overnight. The reaction mixture was diluted with EtOAc (200 mL) and brine (200 mL). The organic was separated and the aqueous phase was further extracted with EtOAc (100 mL). The combined organic (volume ~600 mL) was washed with brine, dried over MgSO_4_ and concentrated under reduced pressure to yield a crude brown liquid (81% purity) that was purified by chromatography on a 120 g ISCO silica gel column (20-40% EtOAc/heptane) to afford the title compound as a yellow foam (15.35g, 77% yield). ^1^H NMR (400 MHz, CDCl_3_)δ: 8.17 (1H, d, J = 8.0 Hz), 8.03 (1H, d, J = 8.0 Hz), 7.94 (2H, dd, J= 1.2, 8.0 Hz), 7.81 (2H, dd, J = 1.2, 8.0 Hz), 7.74 (1H, s), 7.57 – 7.61 (2H, m), 7.43-7.51 (4H, m), 7.31-7.37 (3H, m), 7.24 (1H, dd, J = 4, 8 Hz), 7.07 (1H, dd, J = 4, 8 Hz), 6.94 (1H, d, J = 8 Hz), 6.55 (1H, d, J = 4 Hz), 6.23 (1H, s), 3.71 (1H, d, J = 4 Hz). LCMS m/z: [M-OH]^+^ = 525.53, rt = 1.02 min.

### Ethyl 2-(2-((1H-indol-3-yl)methyl)-1H-indol-3-yl)-2-oxoacetate

To a stirred suspension of lithium aluminum hydride (9.42 g, 248 mmol) in dry 2-methyltetrahydrofuran (250 mL) at −20 °C was added slowly a solution of (1-(phenylsulfonyl)-1H-indol-2-yl)(1-(phenylsulfonyl)-1H-indol-3-yl)methanol (15.35 g, 27.4 mmol) in 100 mL of 2-methyltetrahydrofuran. The flask was covered with aluminum-foil, heated under gentle reflux for 6.5 h and then stirred at room temperature overnight. LCMS revealed the starting material (rt = 1.02 min) was nearly consumed and the product at rt = 0.89 min was a major peak. The reaction mixture was cooled to 0 °C, quenched by adding slowly 9.4 mL of water, 9.4 mL of 15% NaOH and 28.2 mL of water sequentially, stirred for 1 h and filtered through a Celite pad. The filter cake was washed with EtOAc (~500 mL). The filtrate was washed with brine two times, quickly dried over MgSO_4_ and concentrated under reduced pressure to an amber oil that was used without further purification. A solution of the resulting crude 3-((13-((1H-indol-2-yl)methyl)-1H-indole (7.85 g, 25.5 mmol based on 80% purity by LCMS) in dry THF (100mL) was covered with aluminum foil and cooled to 0 ^o^C. Ethyl oxalyl chloride (2.70 mL, 24.1 mmol) was dissolved in 25 mL of THF and added dropwise over a period of 60 min. The mixture was stirred at room temperature overnight and then partitioned between CH_2_Cl_2_ (250 mL) and 5% sodium bicarbonate (100 mL). The organic layer was separated, washed with brine, dried over MgSO_4_ and concentrated under reduced pressure. The residue was recrystallized in acetonitrile (~75 mL) to afford the title compound as a pale yellow solid (3.65 g). The filtrate was concentrated to half volume to provide a second batch (0.95 g). Total = 4.60 g, 52% yield. ^1^H NMR (400 MHz, CDCl_3_) δ : 8.44 (1H, br s), 8.28 (1H, br s), 7.97 (1H, d, J= 8.0 Hz), 7.50 (2H, d, J = 8.5 Hz), 7.24 - 7.33 (3H, m), 7.13 - 7.23 (3H, m), 4.69 (2H, s), 4.51 (2H, q, J = 7.3 Hz), 1.46 (3H, t, J = 7.2 Hz). LCMS m/z: [M+H]^+^ = 347.34, rt = 0.85 min.

### Ethyl 5,11-dihydroindolo[3,2-b]carbazole-6-carboxylate

Into a solution of ethyl 2-(2-((1H-indol-3-yl)methyl)-1H-indol-3-yl)-2-oxoacetate (4.99 g,14.4 mmol in 1,4-dioxane (150 mL) was added methanesulfonic acid (7.5 mL, 115 mmol) slowly. The resulting mixture was heated at 105 °C under gentle reflux for 30 min. LCMS revealed complete conversion to a single product. The mixture was cooled and concentrated under reduced pressure to a minimal volume. The residue was suspended in 5% NaHCO_3_ (250 mL) and stirred for 1 h. The solid was collected by filtration, washed with EtOAc and water, and dried in air flow to afford the title compound as a yellow solid (3.72 g). The organic was separated from the filtrate, washed with brine, dried over MgSO_4_ and concentrated under reduced pressure to give a second batch of the title compound (1.0 g). Total = 4.72 g, 99 % yield. mp 175-177°C (decomp.). ^1^H NMR (400 MHz, DMSO-*d*_6_) δ : 11.51 (1H, s), 10.95 (1H, s), 8.72 (1H, d, J= 8.3 Hz), 8.49 (1H, s), 8.28 (1H, d, J = 7.8 Hz), 7.69 (1H, d, J = 8.0 Hz), 7.54 (1H, d, J = 8Hz), 7.45 (2H, m), 7.19 (2H, m), 4.69 (2H, q, J = 7.2Hz), 1.53 (3H, t, J = 7.2 Hz). ^13^C NMR (100 MHz, DMSO-d6) δ : 167.5, 142.1, 141.6, 136.2, 135.6, 126.7, 126.6, 125.4, 123.8, 122.1, 121.7, 120.8, 120.5, 119.1, 118.2, 112.0, 111.2, 107.2, 105.6, 61.5, 14.5. LCMS m/z: [M+H]^+^ = 329.52, rt = 0.99 min.

### (5,11-Dihydroindolo[3,2-b]carbazol-6-yl)methanol

Into a solution of ethyl 5,11-dihydroindolo[3,2-b]carbazole-6-carboxylate (1.12 g, 3.41 mmol) in THF (75 mL) was slowly added borane-methyl sulfide complex in THF (5.2 mL, 10.4 mmol) at 0 °C. The resulting homogenous solution was heated under reflux overnight. LCMS showed complete conversion to a single product. The reaction mixture was cooled to 0 °C, slowly quenched with water and then diluted with EtOAc. The organic layer was washed with brine, dried over MgSO_4_ and concentrated under reduced pressure. The residue was suspended in hot EtOAc (20 mL), sonicated, cooled to room temperature and filtered to afford the title compound as a light yellow solid (0.63 g). The filtrate was concentrated, taken up in EtOAc (15 mL), sonicated, heated to boiling, cooled to room temperature, filtered and washed with 50% EtOAc/heptane to give a second batch of the title compound (0.26 g). Total = 0.89 g, 89% yield. mp >225 °C. ^1^H NMR (400 MHz, DMSO-*d*_6_) δ : 11.07 (1H, s), 10.94 (1H, s), 8.30 (1H, d, J= 7.8 Hz), 8.19 (1H, d, J = 7.8Hz), 8.06 (1H, s), 7.75 (2H, dd, J = 18.9, 7.8 Hz), 7.38 (2H, m), 7.13 (2H, q, J = 7.3Hz), 5.43 (2H, d, J = 5.0Hz), 5.33 (1H, t, J = 5.0Hz). ^13^C NMR (100 MHz,DMSO-*d*_6_) δ : 141.6, 141.6, 135.8, 134.3, 125.9, 125.4, 123.8, 123.0, 122.9, 122.9, 121.2, 120.6, 118.1, 118.0, 117.0, 111.1, 110.6, 100.1, 58.2. LCMS m/z: [M+H]^+^ = 287.29, rt = 0.78 min.

### 6-Formylindolo[3,2-b]carbazole (FICZ)

Into a solution of (5,11-dihydroindolo[3,2-b]carbazol-6-yl)methanol (0.63 g, 2.20 mmol) in 1,4-dioxane (50 mL) was added DDQ (0.55 g, 2.42 mmol). The resulting mixture was stirred at room temperature overnight. The solid was collected by filtration, washed with dioxane, suspended in water (125 mL) and saturated sodium bicarbonate (75 mL) and stirred at room temperature for 2 h. The mixture was sonicated to break up any large blocks of solid. The solid was collected by filtration, washed with water and dried in a vacuum oven (~60 °C) to afford the first batch of FICZ as a yellow solid (0.39 g). The filtrate from the first collection was concentrated under reduced pressure, partitioned between EtOAc and 5% NaHCO_3_ and stirred overnight. The organic layer was separated and the aqueous was extracted with EtOAc two times. The combined organic was washed with 5% NaHCO_3_ and brine, dried over MgSO_4_ and concentrated. The residue was suspended in hot EtOAc, sonicated, cooled to room temperature, filtered, washed with fresh EtOAc and dried under reduced pressure to obtain a second batch of FICZ (0.18 g). Total yield = 0.57 g, 90 %. mp >225°C. ^1^H NMR (400 MHz, DMSO-*d*_6_) δ :11.75 (1H, s), 11.63 (1H, s), 11.37 (1H, s), 8.61 (1H, s), 8.57 (1H, d, J= 8.0 Hz), 8.31 (1H, d, J = 7.5Hz), 7.75 (1H, d, J = 8Hz), 7.61 (1H, d, J = 8.3 Hz), 7.47 (2H, dt, J = 15.6, 7.5, 7.5 Hz), 7.22 (2H, m). ^13^C NMR (100 MHz, DMSO-*d*_6_) δ : 190.5, 142.1, 142.1, 135.8, 135.2, 126.9, 126.7, 125.1, 123.8, 122.1, 121.8, 121.5, 120.9, 119.7, 119.3, 112.8, 112.5, 111.9, 110.4. LCMS m/z: [M+H]^+^ = 285.25, rt = 0.91 min. HRMS calcd for C19H12N2O: 284.0950; Found: 284.0954.

### Indolo[3,2-b]carbazole-6,12-dione

FICZ (80 mg, 0.281 mmol) was suspended in MeOH (15 mL) in a 500 mL Erlenmeyer flask and covered with plastic film to prevent evaporation. The flask was irradiated in a chamber with illuminance at 11000 lux with occasional shaking. LCMS indicated very slow decomposition due to the low solubility of FICZ in MeOH. After 10 days, LCMS revealed no residual FICZ was present. The suspension was concentrated under reduced pressure. The residue was suspended in CHCl_3_, stirred and filtered. The residue that was insoluble in CHCl_3_ was collected, washed with CHCl_3_, and dried to yield the title compound as a yellow solid (40 mg, 47 %). ^1^H NMR (400 MHz, DMSO-d_6_) δ: 12.92 (2H, s), 8.07 (2H, d, J= 6.8 Hz), 7.54 (2H, d, J = 7.3 Hz), 7.38-7.28 (4H, m). ^13^C NMR (100 MHz, DMSO- d_6_) δ: 176.3, 140.1, 137.9, 125.9, 124.5, 124.4, 121.8, 115.3, 114.5. LCMS m/z: [M+H]^+^ = 287.16, rt = 0.80 min.

### Photostability test

Method 1: FICZ (15.0 mg) and maleic acid (internal NMR standard, 6.0 mg) were dissolved in 5 mL of DMSO-*d*_6_. The solution was analyzed by ^1^H NMR as time point zero. 0.5 mL of the solution was placed into separate ICH 20 mL Flint clear glass bottles that were irradiated in a photostability chamber with illuminance set to 11000 lux as measured by a digital light meter. The contents of separate bottles were checked by ^1^H NMR at 1 h, 2 h, 3 h, 4 h, 5 h, 7 h, 10 h, 16 h, and 24 h. Residual FICZ was quantified by the peak height of the CHO proton (11.37 ppm) compared to the height of the vinyl proton of maleic acid (6.27 ppm). Decomposition of FICZ was also monitored by LCMS as additional confirmation of the NMR quantification. A parallel experiment was run under the same conditions with indolo[3,2-b]carbazole (12.4 mg) and maleic acid (6.4 mg) in 5.1 mL of DMSO-*d*_6_ and stability data collected at 1 h, 2 h, 4 h, 6 h, 12 h, 18 h, and 24 h.

Method 2: FICZ (8.5 mg) and maleic acid (3.5 mg) were dissolved in 4 mL of DMSO-*d*_6_. The solution was analyzed by ^1^H NMR as time point zero. 0.5 mL of the solution was placed into separate ICH 20 mL Flint clear glass bottles that were placed in direct sunlight with illuminance of 63000 lux as measured by a digital light meter (from 1:00PM to 3:40PM on March 17, 2015, a sunny day with minimal clouds in Durham, NC, USA). The contents of separate bottles were checked by ^1^H NMR at 1.5 min, 3 min, 10 min, 30 min, and 93 min. Residual FICZ was quantified by the peak height of the CHO proton (11.37 ppm) compared to the height of the vinyl proton of maleic acid (6.27 ppm).

Method 3: FICZ (14.4 mg) and maleic acid (internal NMR standard, 6.4 mg) were dissolved in 5 mL of DMSO-*d*_6_. The solution was analyzed by NMR as time point zero. The solution was then divided into four ICH 20ml Flint clear glass bottles: Bottle A was exposed to air and was unprotected from the bright light; Bottle B: was flushed with nitrogen and left unprotected from the bright light; Bottle C was exposed to air but covered by aluminum-foil; and Bottle D was flushed with nitrogen and covered by aluminum foil. All bottles were placed in a photostability chamber with illuminance set to 11100 lux. The contents of each bottle were checked by ^1^H NMR at 1 d, 4 d, 8 d, 14 d and 21 d using the quantification procedure described in Method 1.

### Biological assay

CYP1A1-bla-LS180 cells (Invitrogen) were suspended in assay media (OptiMEM, 1% charcoal stripped serum, 1 mM sodium pyruvate, 0.1 mM NEAA) at a density of 20000 cells/mL. The compounds were dissolved in DMSO at a concentration of 10 mM. 0.5 mL of the compound solutions were added to a 384 well black, clear bottom, cell-coat assay plate (Griener Bio-one). A serial dilution at a ratio of 1:3 was made across the plate using assay media. A 50 μL aliquot of cell suspension was added to each well, and the compound plate was incubated overnight at 37 °C in 5 % CO_2_. The plate was equilibrated at room temperature for 30 min before adding 10 μL of the LiveBLAzer^TM^ FRET B/G substrate solution (Life Technologies) to each well. The plate was incubated for 90 min at room temperature in the dark before being read on a Perkin Elmer Envision using the bottom read fluorescence program with excitation at 400 nm and detection at 460/535 nm. Results for each test well were expressed as % inhibition using DMSO as the 0% control and 1 μM FICZ as the 100% control.

## Results

### Practical synthesis of FICZ

The readily available 1-(phenylsulfonyl)-1H-indole (**2**, Fig 2) was selected as one of the starting materials, since the phenylsufonyl group protecting group would direct ortho-lithiation of the indole by *n*-BuLi [14]. Reaction of the 2-lithiate salt **3** with commercially available 1-(phenylsulfonyl)-1H-indole-3-carbaldehyde (**4**) afforded the desired bis-indole intermediate **5** in a good yield following chromatography.

**Fig 2.**
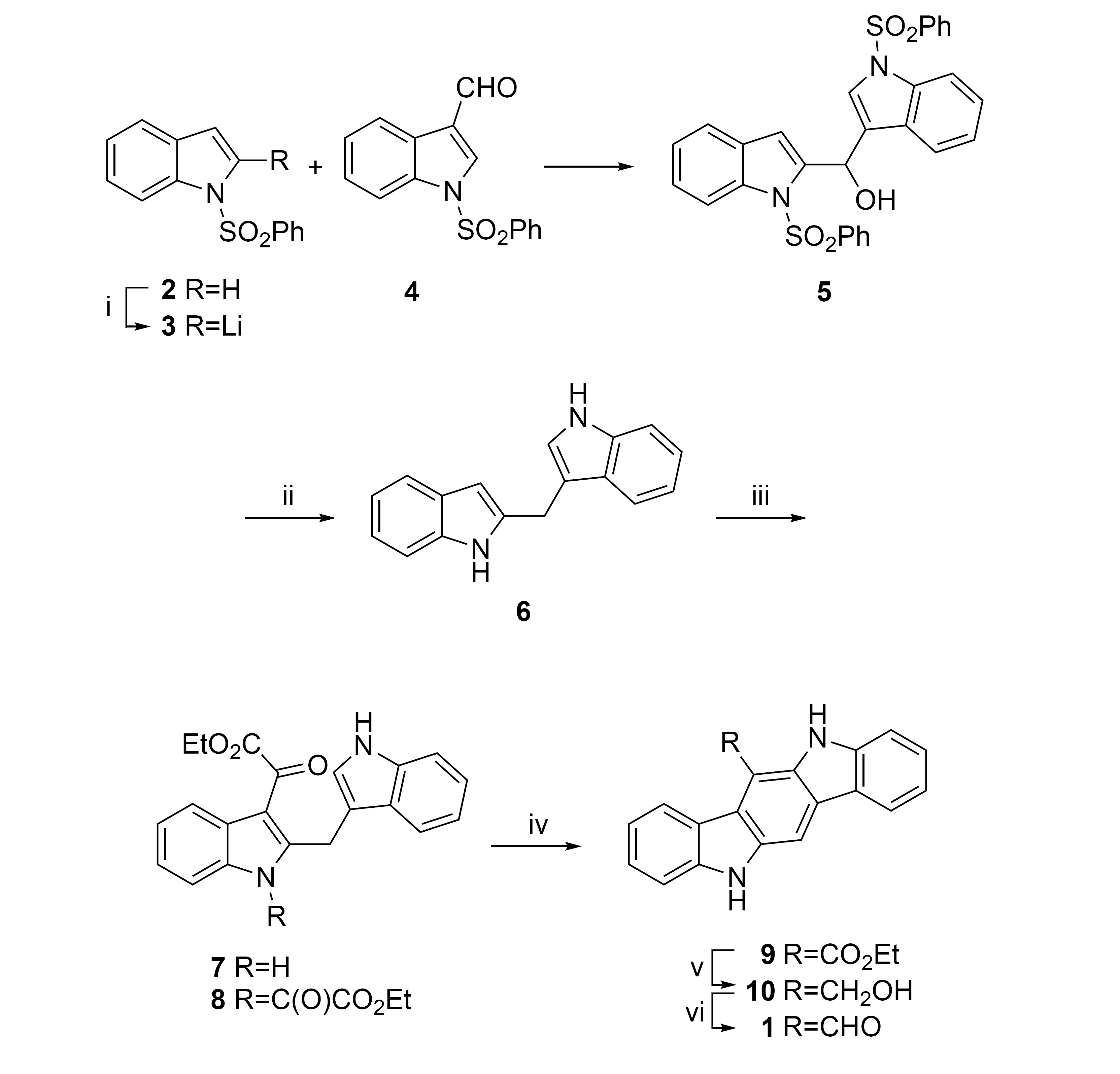
Practical chemical synthesis of FICZ (1). (i) *n*-BuLi, THF, −70 °C. 77%. (ii) LiAlH_4_, THF, reflux. (iii) EtO_2_CCOCl, pyridine, THF. 52% for two steps. (iv) MeSO_3_H, dioxane, reflux. 99%. (v) BH_3_•Me_2_S, THF, reflux. 89%. (vi) DDQ, dioxane. 90%.

Removal of the phenylsulfonyl groups and benzylic hydroxyl was achieved by heating with excess lithium aluminium hydride followed by a standard Fieser work up protocol to afford crude **6** at 80% purity. The limited stability of **6** [10] required use of the crude material without further purification. Reaction of **6** with excess ethyl oxalyl chloride did not yield the desired 3-acylated indole 7, but instead produced the double-acylated product **8** due to the NH group reacting as a vinylogous amide. Fortunately, reaction of crude 6 with 0.95 equivalents of ethyl oxalyl chloride yielded pure mono-carboxy 7 in good yield following crystallization from acetonitrile. Ring closure and carbazole formation was achieved by heating 7 in dioxane with methanesulfonic acid followed by work up and crystallization to deliver pure carbazole ester **9**. Borane-methyl sulfide complex was found to be the optimal reagent for reduction of the ester **9** due to the homogenous reaction mixture and simple work up to produce alcohol **10**. Oxidation of 10 by DDQ yielded aldehyde **1**, which was isolated by filtration. Following extensive washing with water, analytically pure FICZ (1) was produced as a yellow solid in high yield. The synthetic procedure (Fig 2) was successfully run on a multi-gram scale. Purification of the ring-closed carbazoles by crystallization was critical, since their poor solubility made column chromatography on large scale impractical.

### Photostability of FICZ

Although FICZ (**1**) was stable as a pure solid, we observed decomposition of DMSO solutions upon standing under laboratory lighting. To measure the photostability of 1 by NMR, a deuterated DMSO solution was placed in a clear Flint bottle inside a closed chamber and subjected to a light intensity of 11000 lux (approximately equivalent to daylight without direct sun exposure). Maleic acid was employed as an internal NMR reference. The unsubstituted indolo[3,2-b]carbazole (**11**, Fig 3a), a chemically-related AhR ligand, was included as a comparator. As shown in Fig 3b, **1** decomposed rapidly in the presence of air and light with a half-life of less than 3 h. Decomposition of the unsubstituted carbazole **11** was at least 10-times slower under identical conditions. When the experiment was repeated in direct sunlight at a light intensity of 63000 lux, decomposition of **1** was even more rapid with a half-life of ~30 min (Fig 3c). HPLC analysis indicated that both **1** and **11** formed a single major decomposition product, which was isolated and identified as indolo[3,2-b]carbazole-6,12-dione (12) by MS, ^1^H and ^13^C NMR (Fig 3a) [15]. To further characterize the stability of **1**, DMSO solutions were subjected to 11000 lux over 28 d under different storage conditions (Fig 3d). Significant decomposition was observed under all conditions, however the rate varied depending on the presence of bright light or air. Replicating the prior conditions of bright light and air, 1 was completely decomposed within 1 day (Condition A). When the bottle was flushed with nitrogen to minimize the presence of air, decomposition of **1** was much slower (Condition B). In contrast, protection of the bottle from light exposure with aluminum foil further slowed, but did not prevent the decomposition (Condition C). Finally, removal of air and protection from light (Condition D) yielded the slowest rate of decomposition. In summary, **1** showed a wide range of stability as a solution in DMSO. Decomposition required the presence air and was greatly accelerated by bright light. Importantly, we observed that air or light alone was sufficient to promote significant decomposition of FICZ (**1**) as a DMSO solution upon prolonged storage.

**Fig 3.**
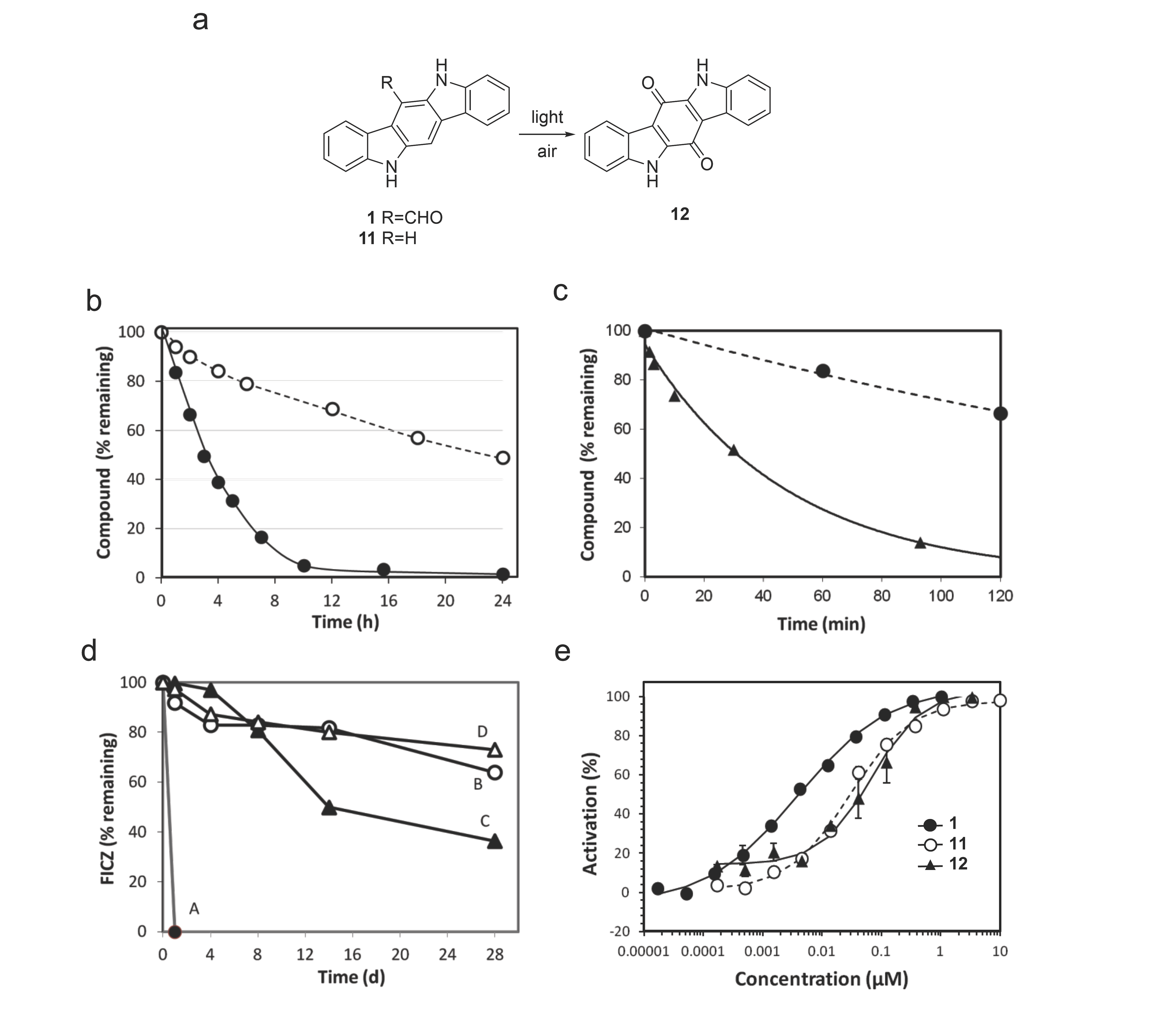
FICZ photostability. a) FICZ **(1)**and indolo[3,2-b]carbazole **(11)**decompose to indolo[3,2-b]carbazole-6,12-dione **(12);**b) Time course of FICZ **(1)**● and indolo[3,2-bjcarbazole (11) ○ stability in air and bright light; c) Time course of FICZ (1) stability in air and bright light ● or direct sunlight ▲; d) FICZ (1) stability in (A) air and light ●, (B) nitrogen and light ○, (C) air and dark ▲, and (D) nitrogen and dark Δ; e) Activation of the human AhR by FICZ **(1),**indolo[3,2-b]carbazole **(11),**and indolo[3,2-b]carbazole-6,12-dione **(12),**n=2 for all data.

### Bioactivity of FICZ and its photoproduct

To understand the biological consequences of the decomposition of **1**, a freshly prepared solution was tested for its ability to activate the human AhR. Using cells that were engineered to express a reporter gene composed of the *cyp1a1* promoter driving expression of a β-lactamase reporter (Fig 3e), **1** activated AhR with an EC_50_ value of 3.9 nM. Under identical conditions, the photo-decomposition product **12** showed a much lower EC_50_ value of 70 nM, while the unsubstituted carbazole **11** had an EC_50_ value of 30 nM. Thus, photo-decomposition of FICZ (1) to the quinone (**12**) would result in a large decrease in potency (>17-fold) for the activation of AhR, a difference that could to be biologically significant for an endogenous hormone [16]. In contrast, conversion of unsubstituted carbazole (**11**) to the quinone (**12**) led to only a modest 2-fold decrease in potency that is unlikely to be biologically significant.

## Discussion

FICZ (**1**) demonstrates many of the properties of a light-dependent hormone that could entrain peripheral tissues to the day/night cycle through exposure to sunlight. 1 is formed by UV or light irradiation of tryptophan [17] and, following activation of AhR, can increase expression of CYP1A1 to induce its own metabolism [7, 18] and provide a feedback mechanism to eliminate the hormone. Our stability experiments now reveal an alternative pathway to limit exposure of peripheral tissues to high levels of FICZ (**1**), through a non-enzymatic light and air-induced oxidation of the formyl-substituted carbazole. Notably, photo-decomposition of **1** was much more rapid than the unsubstituted analog **11**, an alternate AhR ligand that can be formed in the gut from dietary precursors but is not induced by light. Although, both carbazoles decomposed to the identical quinone **12**, the presence of the aldehyde in **1** clearly accelerated photo-induced oxidation of the carbazole core. The stability experiments were conducted using light intensities comparable to normal daylight as well as in direct bright sunlight. We observed that decomposition was faster in the brighter light corresponding to direct sun exposure. Although regular laboratory lighting is generally 20-100 times less intense than daylight, our results dictate that solutions of **1** should be prepared immediately prior to biological testing and be shielded from air and light to ensure accurate results.

FICZ (**1**) is a naturally-occurring AhR ligand that is formed in the skin [6, 19] and may be an important physiological regulator of this transcription factor. Many dermatological conditions, such as atopic dermatitis and psoriasis, can be alleviate though judicious sun exposure [20]. In fact, therapeutic UV-B dosing is a common treatment for both conditions. FICZ (**1**) has demonstrated immune modulating effects through activation of AhR and is effective in treating animal models of psoriasis [21]. Pharmacological treatment of atopic dermatitis and psoriasis with topical AhR ligands has also been shown to be effective in human clinical trials [22, 23]. Thus, the endogenous light-regulated level of FICZ may have an important role in maintaining healthy skin [24]. In addition to the production of **1** by the action of light on the skin, our results now suggest the possibility that the combination of light exposure with free oxygen in cells provides an additional non-enzymatic mechanism for limiting the total levels of FICZ (**1**) in peripheral tissues.

## Conclusion

We have developed the first efficient gram scale synthesis of FICZ (**1**), which will allow investigators ready access to this important tool for study of AhR biology in pharmacological models of disease. Our results provide support for the proposed role of FICZ (**1**) as a light-dependent switch for a ligand-dependent transcription factor in peripheral tissues, such as the skin. It is remarkable that nature has non-enzymatic light-facilitated mechanisms to both generate [4, 5] and to remove this signaling molecule. Whether this process functions in humans to limit the concentration of FICZ in skin under conditions of bright sunlight remains to be explored, but it could potentially add further support to its emerging role as a physiologically relevant light-dependent hormone.

## Acknowledgment

The SGC is a registered charity (number 1097737) that receives funds from AbbVie, Bayer Pharma AG, Boehringer Ingelheim, Canada Foundation for Innovation, Eshelman Institute for Innovation, Genome Canada, Innovative Medicines Initiative (EU/EFPIA), Janssen, Merck & Co., Novartis Pharma AG, Ontario Ministry of Economic Development and Innovation, Pfizer, São Paulo Research Foundation-FAPESP, Takeda, and Wellcome Trust.

